# Potential for virus endogenization in humans through testicular germ cell infection: the case of HIV

**DOI:** 10.1101/2020.06.04.135657

**Authors:** Dominique Mahé, Giulia Matusali, Claire Deleage, Raquel L. L. S. Alvarenga, Anne-Pascale Satie, Amélie Pagliuzza, Romain Mathieu, Sylvain Lavoué, Bernard Jégou, Luiz R. de França, Nicolas Chomont, Laurent Houzet, Antoine D. Rolland, Nathalie Dejucq-Rainsford

**Affiliations:** Univ Rennes, Inserm, EHESP, Irset (Institut de recherche en santé, environnement et travail) - UMR_S1085, F-35000 Rennes, France; AIDS and Cancer Virus Program, Leidos Biomedical Research, Inc. Frederick National Laboratory for Cancer Research, Frederick, Maryland, USA; Laboratory of Cellular Biology, Department of Morphology, Federal University of Minas Gerais, Belo Horizonte, Brazil; Department of Microbiology, Infectiology and Immunology, Faculty of Medecine, Université de Montréal, and Centre de recherche du CHUM, Montréal, Quebec, Canada; Centre Hospitalier Universitaire de Pontchaillou, Service Urologie, Rennes, France; Centre Hospitalier Universitaire de Pontchaillou, Centre de Coordination des prélèvements, Rennes, France

**Author notes:** Corresponding author: Nathalie DEJUCQ-RAINSFORD, IRSET-Inserm U1085, 9 avenue du Pr Léon Bernard, F-35000 Rennes, France.

**Keywords:** virus-HIV, SIV, testis, germ line-spermatogenesis-gametes-entry, integration-replication-endogenization

## Abstract

Viruses have colonized the germ line of our ancestors at several occasions during evolution, leading to the integration in the human genome of viral sequences from over 30 retroviral groups and a few non-retroviruses. Among the recently emerged viruses infecting humans, several target the testis (eg HIV, Zika and Ebola viruses). Here we aimed to investigate whether human testicular germ cells (TGCs) can support integration by HIV, a contemporary retrovirus that started to spread in the human population during the last century. We report that albeit alternative receptors enabled HIV-1 binding to TGCs, HIV virions failed to infect TGCs *in vitro*. Nevertheless, exposure of TGCs to infected lymphocytes, naturally present in the testis from HIV+ men, led to HIV-1 entry, integration and early protein expression. Similarly, cell-associated infection or bypassing viral entry led to HIV-1 integration in a spermatogonial cell line. Using DNAscope, HIV-1 and SIV DNA were detected within a few TGCs in the testis from one infected patient, one rhesus macaque and one African Green monkey *in vivo*. Molecular landscape analysis revealed that early TGCs were enriched in HIV early co-factors up to integration and had overall low antiviral defenses when compared with testicular macrophages and Sertoli cells. In conclusion, our study reveals that TGCs can support the entry and integration of HIV upon cell-associated infection. This could represent a way for this contemporary virus to integrate our germline and become endogenous in the future, as happened during human evolution for a number of viruses.

**Importance:** Viruses have colonized the host germ line at many occasions during evolution to eventually become endogenous. Here we aimed at investigating whether human testicular germ cells (TGCs) can support such viral invasion by studying HIV interactions with TGCs *in vitro*. Our results indicate that isolated primary TGCs express alternative HIV-1 receptors allowing virions binding but not entry. However, HIV-1 entered and integrated in TGCs upon cell-associated infection, and produced low level of viral proteins. *In vivo*, HIV-1 and SIV DNA was detected in a few TGCs. Molecular landscape analysis showed that TGCs have overall weak antiviral defenses. Altogether, our results indicate that human TGCs can support HIV-1 early replication including integration, suggesting potential for endogenization in the future generations.

## Introduction

Retroviruses have repeatedly infected the germline during our evolution and integrated their DNA into the host genome, which has been passed on to the next generations by Mendelian inheritance [1,2]. When not detrimental, the integration of viral sequences has led to their fixation and endogenization in the population [3]. Thus, about 8% of the human genome is now composed of endogenous retroviruses (ERVs), representing 31 distinct viral groups [2]. Such integration, still ongoing in mammals [4], has driven the acquisition of new functions in the host, including a protective role against exogenous infections [5,6]. Within the retrovirus family, several lentiviruses have been endogenized in mammals, including SIV in prolemurians [7–12]. A few non-retroviruses have also colonized the germline of their host but the mechanisms for their integration are unclear [1].

HIV-1 is an emerging zoonotic lentivirus that has infected over 70 million people since its worldwide spreading in the human population from the beginning of the 1980s. Current antiretroviral treatments effectively control but do not eradicate the virus, which still represents a major threat for the human population, with an average of nearly 2 million new infections each year. Horizontal transmission through semen plays a key role for HIV-1 dissemination, and is most likely mediated by the free viral particles and infected leukocytes present in this fluid [13]. We and others found that HIV/SIV strains in semen are locally produced in a subset of individuals (reviewed in [13]) and arise from several organs within the male genital tract [14]. Interestingly, HIV-1 can associate with spermatozoa *in vitro* [15] and was detected by some authors in a small proportion of patients’ sperm [16–25]. Since spermatozoa have a highly compact nucleus and are transcriptionally silent, this detection may suggest infection of their germ cell progenitors in the testis.

The testis is an immune-privilege organ with restricted drugs penetration, considered to constitute a tissue reservoir for HIV and other emerging viruses such as Zika and Ebola [1,26,27]. HIV-1 productively infects testicular T lymphocytes and resident macrophages, which are naturally localized in close proximity to the early germ cells present in the basal compartment of the seminiferous epithelium and therefore outside the blood testis/Sertoli cell barrier [27–31]. A couple of early studies reported HIV nucleic acids in isolated cells resembling germ cells within the seminiferous tubules of deceased patients using *in situ* PCR, a controverted technique suspected of generating false positives [30,32,33]. These findings have therefore been largely dismissed [34,35]. Furthermore, the detection of viral RNA in TGCs could reflect an accumulation of virions bound to the cell surface rather than a true infection. Unfortunately, the scarce access to testicles of HIV-1+ men has hindered further investigations on this debated issue [35]. SIV RNA and proteins were later described in TGCs from non-human primates, using *in situ* hybridization and immunohistochemistry [36,37]. Whether human TGCs are infected by HIV and to which extent remains an open question.

In this study, we aimed to investigate whether human TGCs support HIV entry and integration as a proxy to evaluate potential for viral colonization in the next generations’ human genome. We found that HIV-1 binds to primary TGCs, but that viral entry was inefficient. However, virus integration and early viral protein expression were observed following cell-associated infection. *In vivo*, testicular germ cells harboring viral DNA were detected in the testis from an HIV-infected individual using next generation *in situ* hybridization DNAscope. SIV-infected TGCs were also detected in one experimentally-infected rhesus macaque and one African Green monkey, a natural host for SIV. To determine whether human TGC had evolved elevated defense mechanisms to prevent vertical transmission of viral sequences to the offspring and thus potential endogenization, we analyzed the molecular landscape of TGCs and compared it with that of HIV-permissive and non-permissive testicular somatic cells using single-cell RNA-sequencing data. This analysis revealed relatively low gene expression levels for viral sensors and HIV early life cycle inhibitors in TGCs, together with an enrichment in HIV early co-factors in spermatogonia. Overall, our study provides the proof of concept that human TGCs can support HIV entry and integration, albeit inefficiently. Colonization of the human male germline could therefore lead to the vertical transmission of viral genes and ultimately to their endogenization in the next generations.

## Results

### Isolation and characterization of primary human testicular germ cells (TGCs)

We isolated fresh TGCs from the testes of uninfected donors displaying normal spermatogenesis and characterized them based on their ploidy profile (Fig. 1A, B) as well as expression of specific markers (Fig. 1C, D, E). As expected, three DNA content profiles were present in all TGCs preparations (n=9 donors) with a median value of 54% of n DNA spermatids (range 48-59%), 28% of 2n DNA spermatogonia and secondary spermatocytes (range 22-31%) and 20% of 4n DNA primary spermatocytes and mitotic spermatogonia (range 16-25%) (Fig. 1A, B). The germ cell marker DDX4 was detected in a median of 84% of TGCs (range 70-95%) (Fig. 1C, E). As expected, DDX4 did not label every germ cell because its expression is variable among germ cells, independently of their differentiation stage [38]. TGCs with n, 2n and 4n DNA were DDX4+ with a median value of 80% (72-98%), 61 % (40-95%) and 90% (68-97%), respectively (Fig. 1C). The early germ cell marker MAGEA4, which labels undifferentiated 2n DNA spermatogonia up to 4n DNA pre-leptotene primary spermatocyte [39], was detected in 40% (32-40%) of TGCs (n=3 donors), among which 37% (24-41%) of 2n DNA and 76% (72-86%) of 4n DNA cells (Fig.1C and E). The purity of TGCs preparations was systematically > 94%, as determined using somatic cell markers HLA Class I (median 1%, range 0,4-4%) and vimentin (median 2%, range 1-6%) as well as pan-leukocytes marker CD45 (median 1%, range 0-4%) (Fig. 1D, E).

**Fig. 1.**
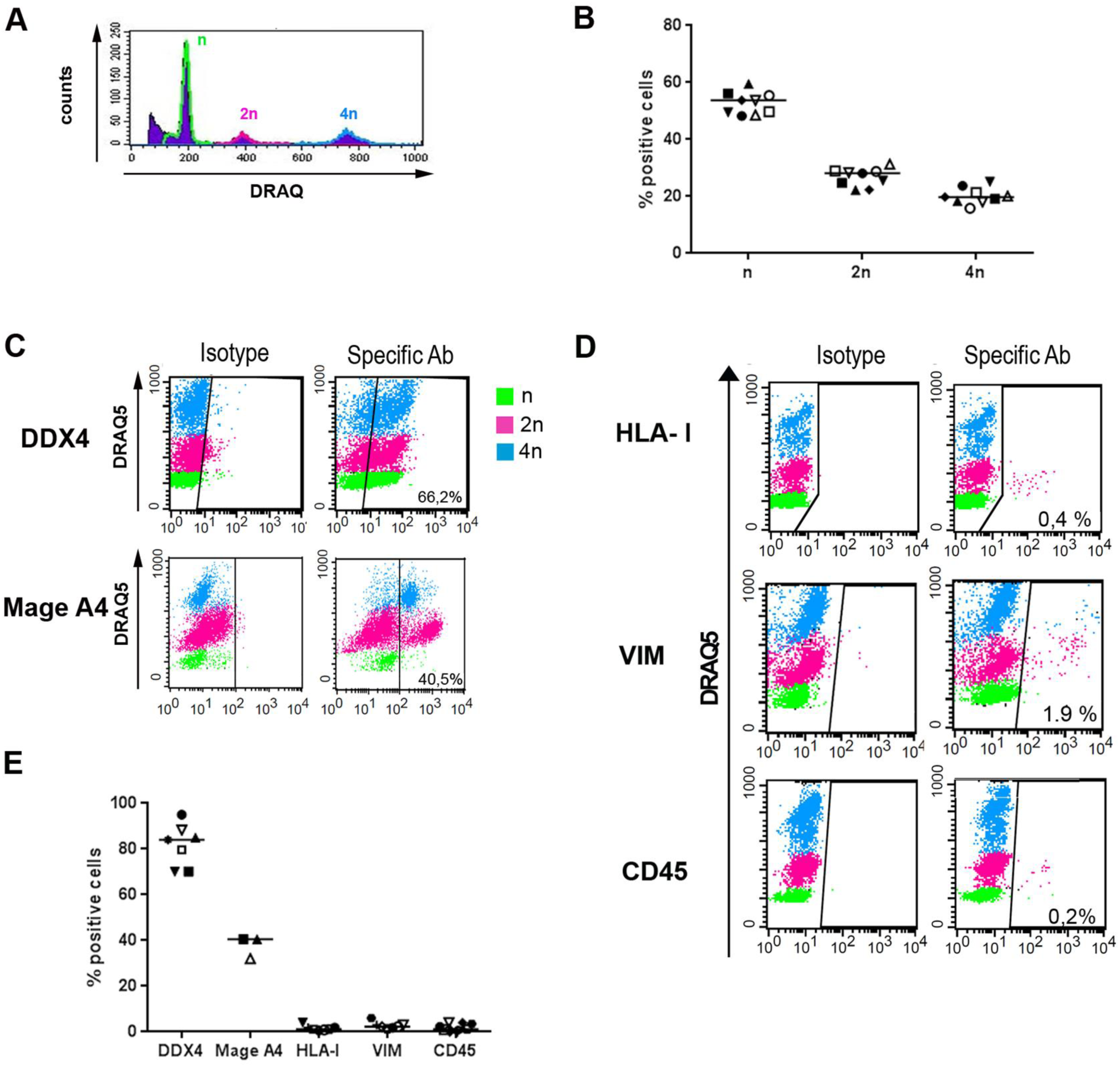
Characterisation and purity of primary TGCs. (**A, B**) DRAQ5 DNA intercalating agent detection in flow cytometry on isolated TGCs showed the presence of haploid (n DNA, green), diploid (2n DNA, pink) and tetraploid cells (4n DNA, blue) (representative profile in A). (**C, E**) As expected, DDX4 labelled most germ cells irrespective of their ploidy, whereas MAGEA4 labelled early diploid and tetraploid germ cells (representative profile in C). (**D, E**) Somatic cell contaminants analysis in isolated TGC populations labeled with DRAQ-5 using flow cytometry and antibodies specific for somatic cells (class I HLA and vimentin) and leukocyte antigen CD45 (representative profile in D).

### HIV binds to TGCs that express alternative receptors on their surface

We first determined whether HIV can attach to TGCs by using the well characterized R5 macrophage-tropic HIV-1_SF162_ (Fig.2) and X4 HIV-1_IIIB_ strains. Dose-dependent binding to TGCs was observed for both strains (Fig.2A and S1). Pronase treatment effectively removed bound HIV-1, indicating that the majority of the attached virions remained at the cell surface (Fig.2A). Similar findings were observed for a primary R5 non-macrophage tropic isolate and a primary R5×4 HIV-1 strain (Fig. 2B). Incubation of TGCs with a protease prior to viral exposure decreased HIV-1 p24 detection in cell lysates by a median of 97% (range 81-100%), indicating that cellular proteins mediated HIV-1 attachment to TGCs (Fig. 2C).

**Fig. 2.**
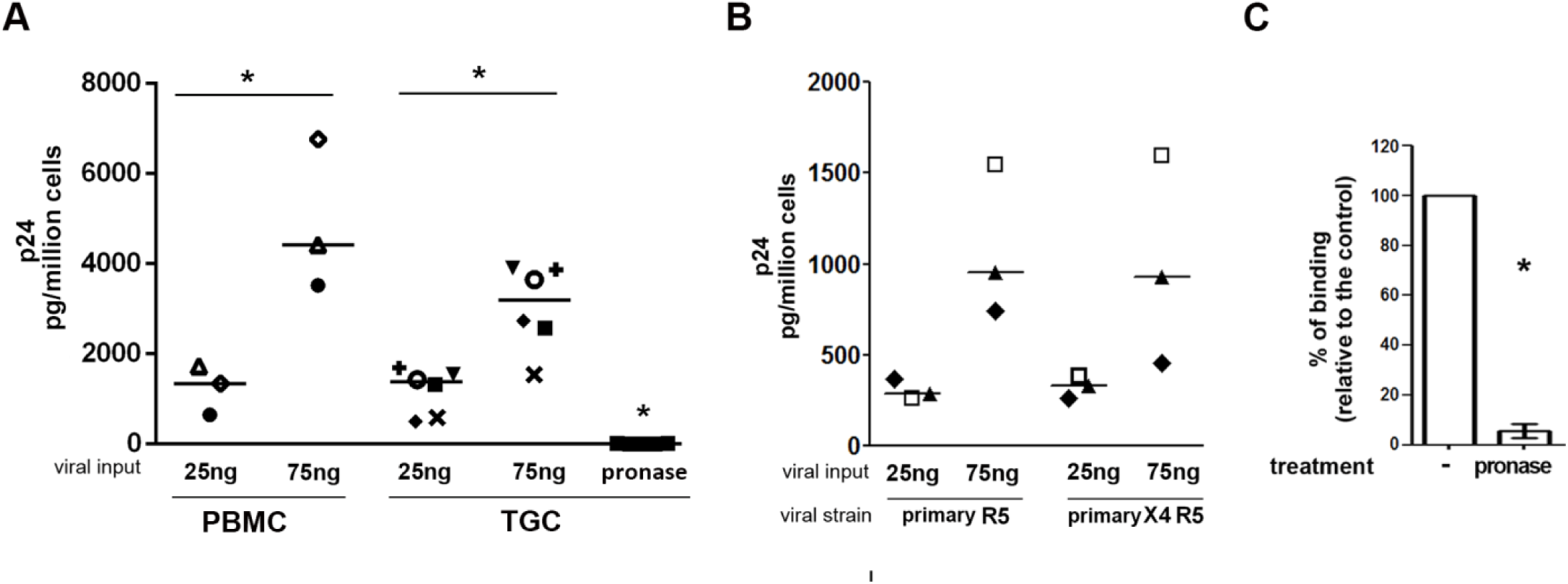
HIV-1 binding to primary TGCs. TGCs isolated from 8 donors or PBMCs used as a positive control were tested for their ability to capture: (**A**) lab-adapted HIV-1 R5_SF162_ or (**B**) primary R5 and R5×4 strains following a 2h incubation at 37°C with the indicated amount of virus, as measured by p24 ELISA on cell lysates following thorough washes and deduction of p24 background measured in wells with no cells. Pronase treatment post-binding was used as a negative control to ensure measure specificity. Statistical analysis with non-parametric test: Wilcoxon test. * p< 0.05; ** p< 0.01; *** p< 0.001. Each donor is represented by a specific symbol. (**C**) Pronase treatment of TGCs prior HIV R5_SF162_ exposure (25ng p24) abrogated viral binding (n=5).

We next explored the expression of candidate receptors for HIV on the cell surface of TGCs (Fig. 3A, B). In agreement with our previous immunohistochemistry data [27], TGCs did not express the main HIV receptor CD4 (Fig. 3A, B). The chemokine HIV co-receptor CCR5 was detected in 4 out of 8 donors on the surface of a very low proportion of TGCs (median 5 %, range 2-12%), whereas CXCR4 was absent in all donors. The chemokine receptor CCR3 was expressed at the TGCs surface in all donors, with a median of 25,9% (14-45%) positive cells (Fig. 3A, B). The alternative HIV binding molecules CD206/mannose receptor, galactosylcerebroside, and heparan sulphate proteoglycans (HSPGs) were detected at the surface of TGCs from all donors, with a median of respectively 74% (51-90%), 77% (51-91%) and 7,1% (3-16%) positive cells (Fig. 3A,B). These receptors were expressed by TGCs irrespective of their DNA content (see representative profiles in Fig. 3A). Competition experiments with BSA-mannose, a ligand for CD206, induced a median reduction of virus attachment of 24% (range 16-29,1%) (Fig. 3C). In contrast, neutralizing antibodies against the cellular glycolipid galactosylceramid had no effect on HIV attachment to TGCs (Fig. 3C), in agreement with the protease experiments. Heparin at 100 U/mL strongly inhibited the capture of HIV particles (median inhibition of 82%, range 79-86%) (Fig. 3C). The contribution of the viral envelope protein gp120 to binding was assessed using *env*-deleted pseudoviruses and antibodies against gp120 (Fig. 3D, E). In 4 independent experiments, the capture of HIV-1 by TGCs was reduced by a median of 53,4% (range 52,6-54,7%) in the absence of *env* (Fig. 3D), while Gp120 neutralizing antibodies reduced HIV binding to TGCs by a median of 29% (range 17-39%) (Fig. 3E), indicating that specific interactions of gp120 with cell surface receptors favored attachment.

**Fig. 3.**
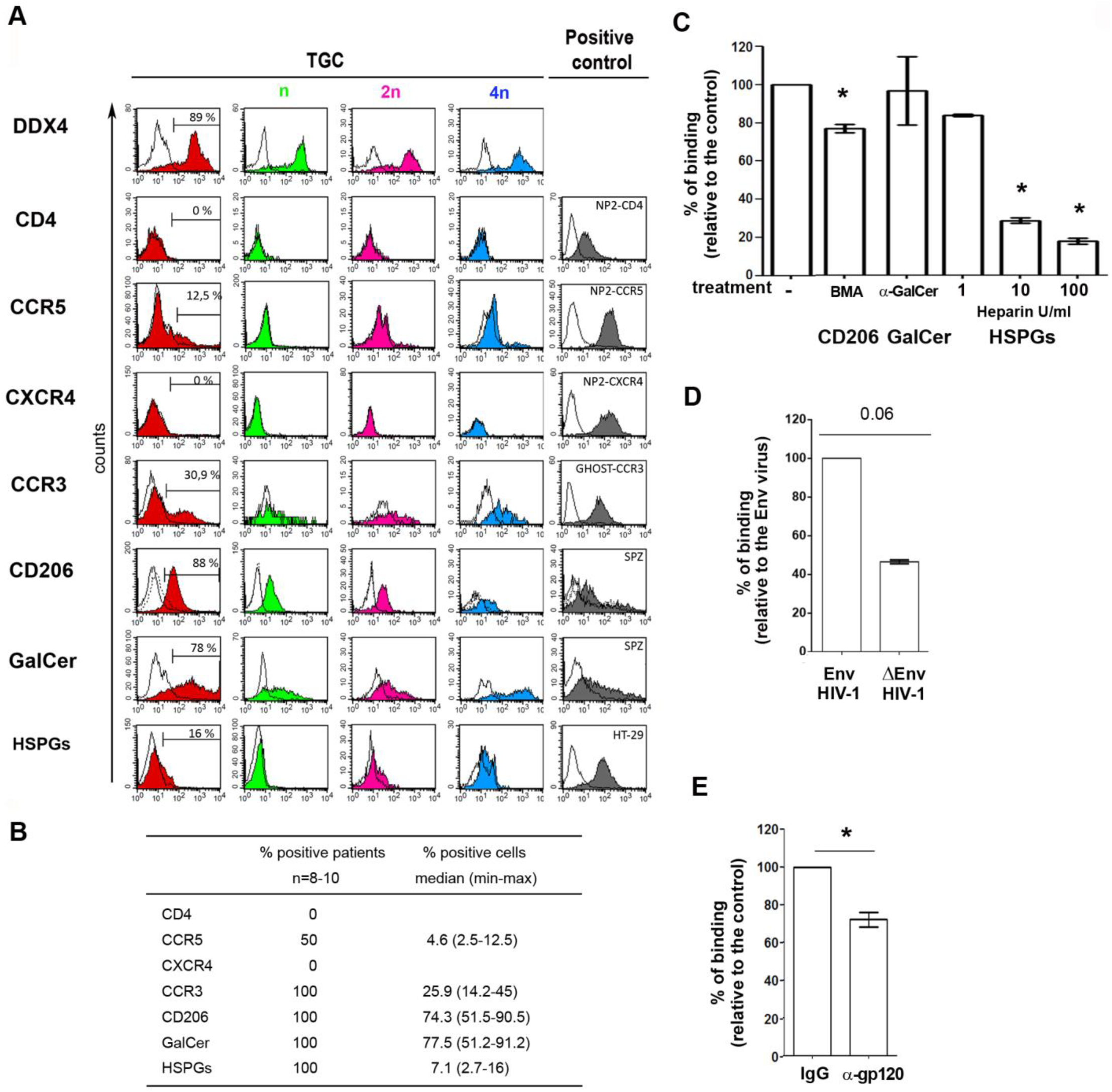
HIV-1 alternative receptors expression by primary TGCs and HIV-1 binding inhibition. (**A**) Representative profiles of membrane expression of HIV receptors on the whole population of TGC (red) and the n (green), 2n (pink) and 4n DNA (blue) TGC populations. Immunolabeling was performed using specific antibodies against CD4, CCR5, CXCR4, CCR3, GalCer or HSPGs, or, in the case of CD206, after detection of biotinylated-BSA-Mannose (BMA) ligand in the presence (dot-lined histogram) or absence (filled histogram) of mannan competitor. BSA was used as control (open histogram). Cells were further stained with DRAQ-5 for ploidy profile. Specific antibody detection was ensured using isotypes as controls, except for HSPGs where pronase treatment was used. Left panel shows positive controls for each receptor. (**B**) The table shows for each receptor the percentage of positive patients, the median and range of positive cells percentage. (**C**) HIV binding was measured after incubation of TGCs with HIV R5_SF162_ (25ng p24) in the presence or absence of CD206 BSA-mannose competitor (BMA), Galactosylceramide specific Mab, HSPGs competitor (heparin) at the 3 indicated doses. Results represent the mean +/- SEM of 3 to 6 independent experiments performed in duplicate and are expressed relative to their respective controls (virus-cell incubation without receptor inhibitors, or in presence of BSA for BSA-mannose). Statistical analysis with non parametric test : Wilcoxon *: p<0.05, **: p<0.01. (**D**) TGCs obtained from 4 donors were incubated for 2h at 37°C with *env* deleted or HIV-1 env expressing pseudoviruses (25 ng p24) and p24 antigen measured by ELISA. (**E**) Viral env neutralization was performed in the presence of gp-120 specific antibody (n=3). In all experiments, background p24 levels measured in wells without cells was deduced from the values obtained with cells.

### TGCs support HIV-1 entry and integration

We next investigated the ability of HIV to enter TGCs and process post-entry steps of viral replication. A major challenge for the study of primary TGCs *in vitro* is their low survival rate outside of the testis environment, since they require physical contacts and paracrine exchange with feeder somatic cells for their maintenance and development. To overcome this challenge, we set-up a co-culture of primary TGCs with human testicular somatic cells in a specific medium that allow long term maintenance of early germ cells [26]. In this culture system, viability dyes showed over 80% of live TGCs after 12 days. After incubating freshly isolated TGCs with wild type HIV-1 R5 (SF162, BaL, JR-CSF) and X4 (IIIB) strains for 4 hours or overnight, no HIVp24 was detected. Using a fluorescent HIV-1^GFP-Vpr^ isolate, only rare entry events were visualized (Fig. 4A). Because cell-associated infection is much more efficient than cell-free infection and within the testis, early TGCs are close to interstitial lymphocytes and macrophages, we investigated HIV entry and integration in TGCs incubated overnight with HIV-1 infected lymphocytes. Confocal microscopy showed DDX4-positive TGCs next to HIV-1-infected PBLs or Jurkat T cells (Fig. 4B) and revealed viral uptake of HIV-1 p24 by DDX4+ TGCs, some in division (Fig. 4C). Altogether, these results indicate that HIV-1 can enter primary TGCs in a cell-associated manner, whereas in our experimental model cell free infection was not readily detected.

**Fig. 4.**
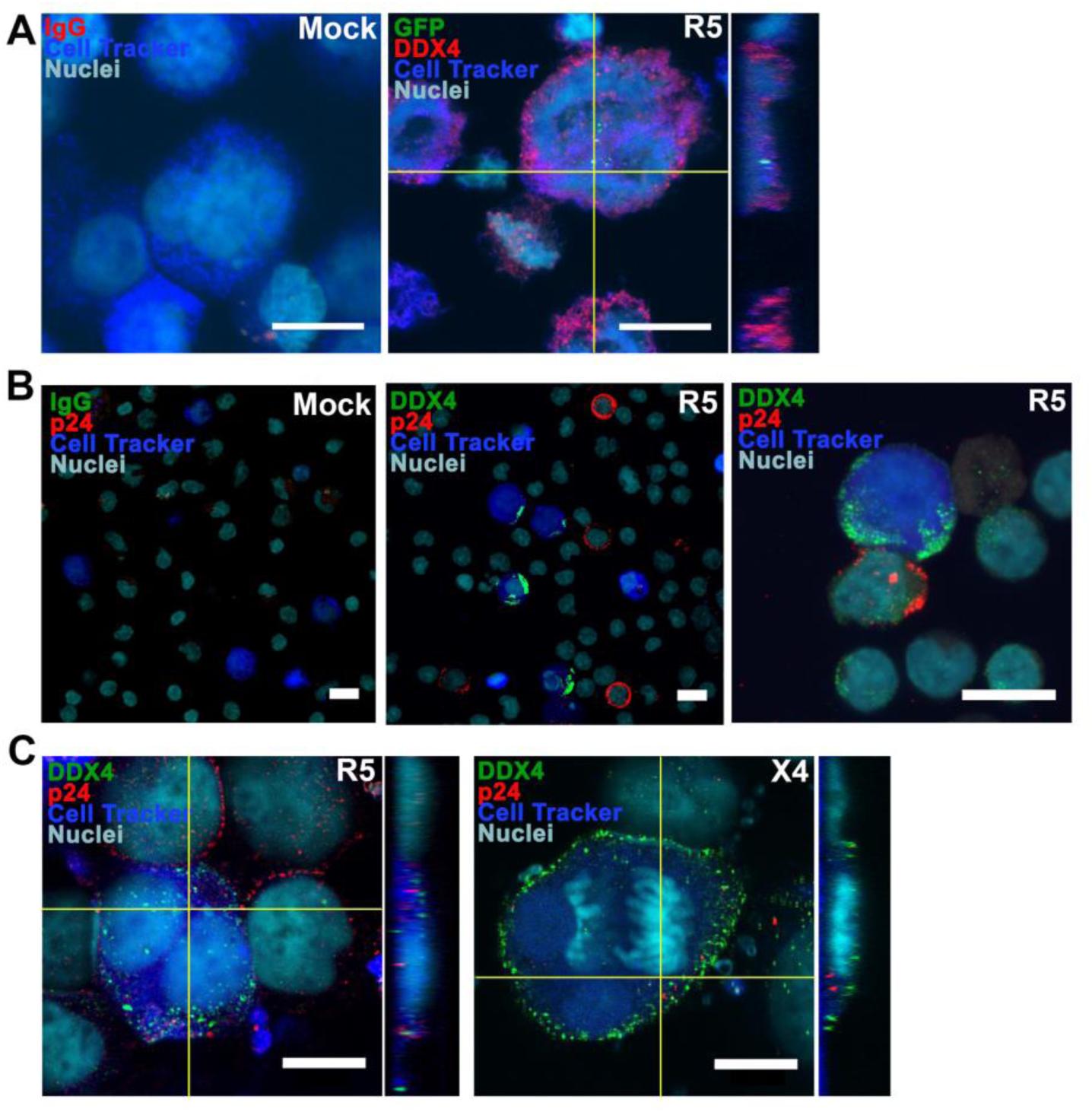
HIV entry in primary TGC following cell-associated infection. (**A**) TGC pre-labelled with CellTracker (blue) were exposed for 18h to R5 HIV-1^GFP-Vpr^ viral particles (green) or mock exposed. Viral entry in cells labelled with DDX4 specific antibody (red) was assessed in confocal microscopy. Mock shows absence of labelling with DDX4 antibody isotype (IgG).. (**B, C**) TGC pre-labelled with CellTracker (blue) were incubated for 18h with either PBMCs infected with a primary R5-tropic HIV-1 strain or mock infected (B), or with Jurkat cells infected with HIV-1 R5 or X4 strains (C). Cells were co-stained for p24 (red) and DDX4 (green) or with control IgG (shown here on mock). (**A, C**) Z-stack and cross-sectional orthogonal viewing are presented to discriminate entering viral particles from membrane bound viruses. Nuclei are stained with DRAQ5 (cyan). Scale bars = 10µm.

To determine whether HIV DNA could integrate into the germ cell genome *in vitro*, freshly isolated TGCs were exposed to infected Jurkat T cells overnight, which were subsequently removed using anti-CD45 magnetic beads. TGCs were cultured for 3 to 7 days and viable cells sorted by flow cytometry, based on DDX4 and MAGEA4 expression and absence of leukocytes and somatic cell markers CD45, CD3 and HLA-DR detection (Fig. 5A, B). As a negative control, TGCs were mixed with infected Jurkat donor cells and were immediately processed (i.e. without incubation and culture) following the same protocol (magnetic beads and FACs sorting) as TGCs incubated ON with donor cells (Fig 5A). Following the cell selection procedure of cultured TGCs, most live germ cells were double positive for DDX4 and MAGEA4 (median 92%, range 82,6-99%), indicating TGCs in the pre-meiotic up to the early meiotic stage. In 3 independent experiments, sorted cultured germ cells were >99% positive for DDX4 and MAGEA4 and negative for immune cell markers CD45, CD3 and HLA-DR (Fig. 5B, C). Alu-gag PCR performed on isolated TGCs demonstrated the presence of integrated HIV DNA in sorted cultured early germ cells from 3 out of 3 donors, although at variable frequencies (Fig. 5C).

**Fig. 5.**
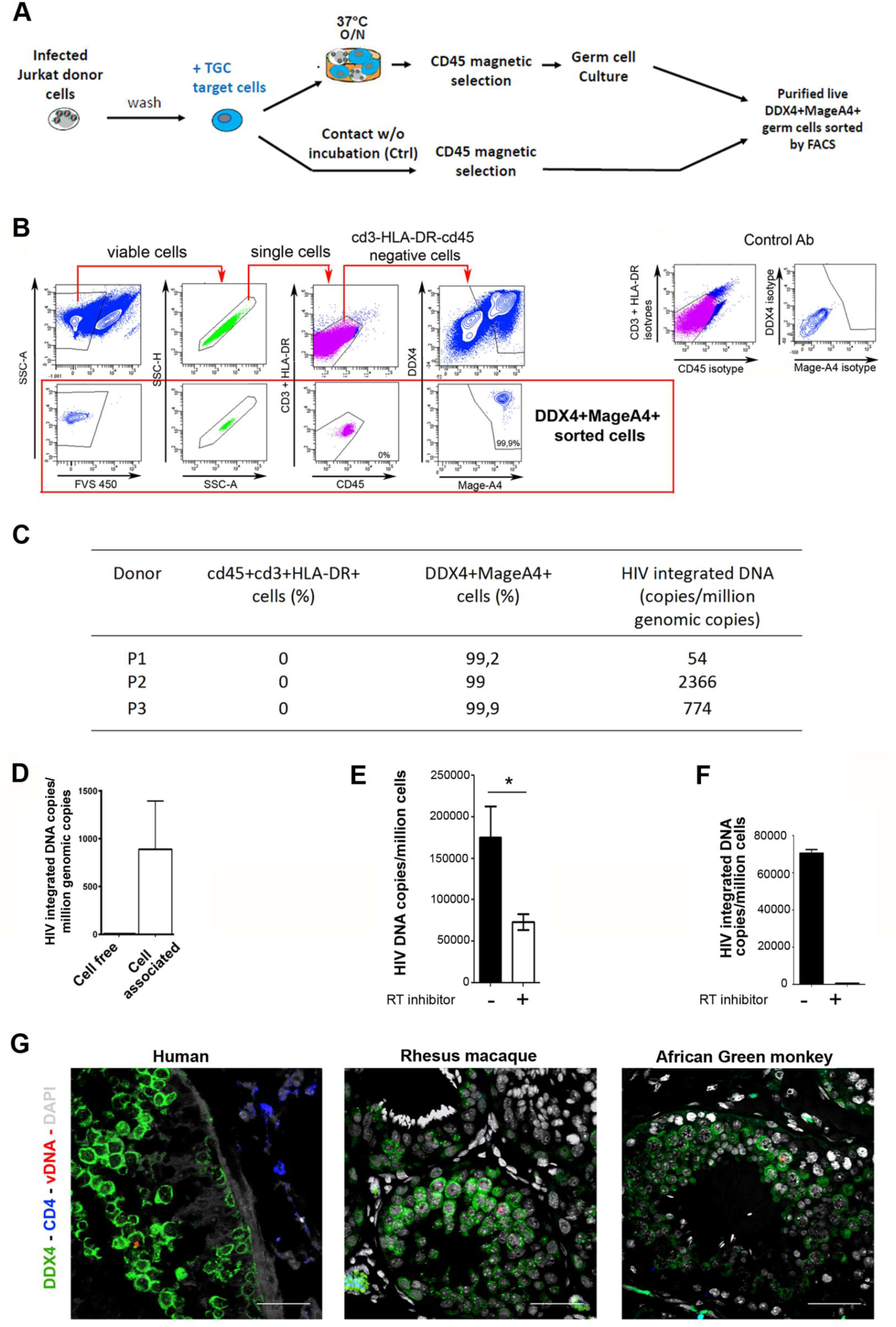
HIV-1 integration in testicular germ cell genome and *in vivo* detection of HIV and SIV DNA in human and non-human primate testicular germ cells. (**A-C**) HIV-1 integration in primary TGCs. (**A**) Experimental design: after overnight contact with Jurkat cells infected with HIV-1 *nef*-ires-GFP, TGCs were recovered following CD45 magnetic selection and put in culture for 5 days. The cells were then submitted to FACS sorting for live cells, negative for leukocytes markers CD45, CD3 and HLA-DR and positive for germ cell markers DDX4 and MAGEA4. As a negative control, TGCs purified by CD45 magnetic selection immediately after contact with Jurkat cells were sorted by FACS similarly to the overnight incubated samples. (**B**) Gating strategy and representative profile of purified live germ cells. Live single cells were selected based on absence of detection of leukocytes markers CD3, HLA-DR and CD45 and on positive expression of DDX4 and MageA4+ germ cell markers. Control antibodies are shown on bottom panel. (**C**) HIV-1 integrated DNA was measured by Alu-gag PCR on 10,000 sorted TGCs exposed ON to Jurkat cells in 3 independent experiments, and the value obtained for the negative controls subtracted. (**D-F**) HIV integration in Tcam-2. (**D**) HIV-1 integrated DNA was measured on Tcam-2 either cultured for 48h following exposure to R5_JR-CSF_ HIV-1 (cell-free) or exposed ON to Jurkat infected with HIV R5_JR-CSF_, purified by CD45 magnetic selection, cultured for 5 days and FACS sorted for CD45 negative live cells (cell associated). (**E, F**) HIV-1 reverse transcription and integration were assessed on Tcam-2 cells exposed to VSV-G-pseudotyped HIV-1. (**E**) Reverse transcripts were detected after 24h culture with or without reverse transcriptase inhibitor nevirapine (n=6) and (**F**) HIV-1 integration assessed by Alu-gag PCR (n=3). (**G**) *In vivo* detection of HIV and SIV DNA in human and non-human primate testicular germ cells by DNAscope. Representative pictures of TGC (green) harbouring viral DNA (red) within testis tissue section from one HIV-1 infected man (VL=51 681 cp/mL), one chronically infected rhesus macaque (VL>10^6 cp/mL) and one African Green monkey 64 days post infection (VL=12085 cp/mL). Scale bars= 100μm.

In order to overcome primary TGCs culture restrictions, we used the well characterized human testicular germ cell line T-cam2, which displays characteristics of stem spermatogonia [40,41]. Except for CXCR4, T-cam 2 cells displayed similar HIV receptor expression than primary TGCs (Fig. S2A) and HIV cell-free entry was also inefficient (Fig. S2B, C). Exposure of T-cam2 cells to Jurkat T cells infected with HIV-1 R5 and X4 strains led to the detection of integrated HIV-1 DNA after flow cytometry sorting of live cells negative for CD45 in 3 independent experiments, whereas no integrated viral DNA was detected after cell-free infection (Fig. 5D). To bypass cell-free HIV entry restrictions, T-cam2 cells were infected with HIV-1 pseudotyped with VSV envelope (VSV G). HIV DNA became readily measurable in T-cam2 cells and its level was significantly decreased by RT inhibitor nevirapine, indicating active reverse-transcription of the viral genome (Fig. 5E). Integrated HIV-1 DNA was detected in T-cam2 cells’ genome exposed to cell free HIV pseudotyped with VSV-G using Alu-gag PCR (Fig.5F). This detection was abrogated in the presence of nevirapine (Fig.5F). Altogether, these results indicate that isolated early germ cells can support low level of HIV-1 DNA integration into their genome.

### HIV-1/SIV DNA is present in TGCs *in vivo*

We next aimed to assess whether TGCs harboring HIV DNA are present *in vivo*. As mentioned, samples of testis tissue from HIV-infected men are extremely rare. Nevertheless we managed to have access to the testis of one HIV+ deceased patient. We screened for HIV DNA+ TGCs using next generation DNAscope *in situ* hybridization for HIV DNA combined with immunofluorescence detection of germ cell marker DDX4. Isolated DDX4+ germ cells harboring viral DNA were found within morphologically normal seminiferous tubules (Fig. 5G). To investigate whether human TGCs are more restrictive to HIV infection than TGCs from non-human primates to SIV, we performed the same approach on the testis from one rhesus macaque and one African green monkey experimentally infected with SIV. In both the rhesus macaque and the African Green monkey, isolated HIV DNA+ TGCs were also observed (Fig. 5G). Overall, these results indicate that human TGCs can support HIV replication up to DNA synthesis *in vivo*, similarly to their non-human primate counterparts, which could represent a way for HIV to become endogenous in the offspring.

### TGCs support low level of HIV replication

Having shown that TGCs can harbour HIV DNA, we aimed to determine whether these cells could support HIV protein expression and virions production. Viral proteins expression in TGCs was analyzed following overnight exposure to Jurkat T cells infected with HIV-1 bearing *nef*-ires-GFP. Nef-GFP was detected using confocal microscopy in primary DDX4+ and MAGEA4+ TGCs after 9 to 12 days of culture (Fig. 6A). Similarly, low level of Nef-GFP was detected by flow cytometry in CD45-negative T-cam2 cells after cell-associated infection, and this detection abolished by pre-treatment of T-cam2 cells with RT-inhibitor nevirapine (Fig. 6B E). Following T-cam2 cells exposure to VSV-G pseudotyped HIV-1-bearing *nef*-ires-GFP, up to 9% of germ cells expressed Nef-GFP at day 5 post-infection, whereas less than 0.2 % of the cells were positive in the presence of nevirapine (Fig. 6C). Inhibitors of both HIV reverse-transcriptase (nevirapine) and integrase (raltegravir) similarly impaired the expression of intracellular p24 (Fig. 6D), demonstrating expression of viral proteins from *de novo* reverse transcribed and integrated virus. The release of infectious particles by T-cam2 cells was confirmed by the infection of permissive human PBLs with T-cam2 supernatants collected at day 5 and day 9 post-infection (Fig. 6E). Altogether, these results show that in addition to integration, early TGCs can support low level of viral production.

**Fig. 6.**
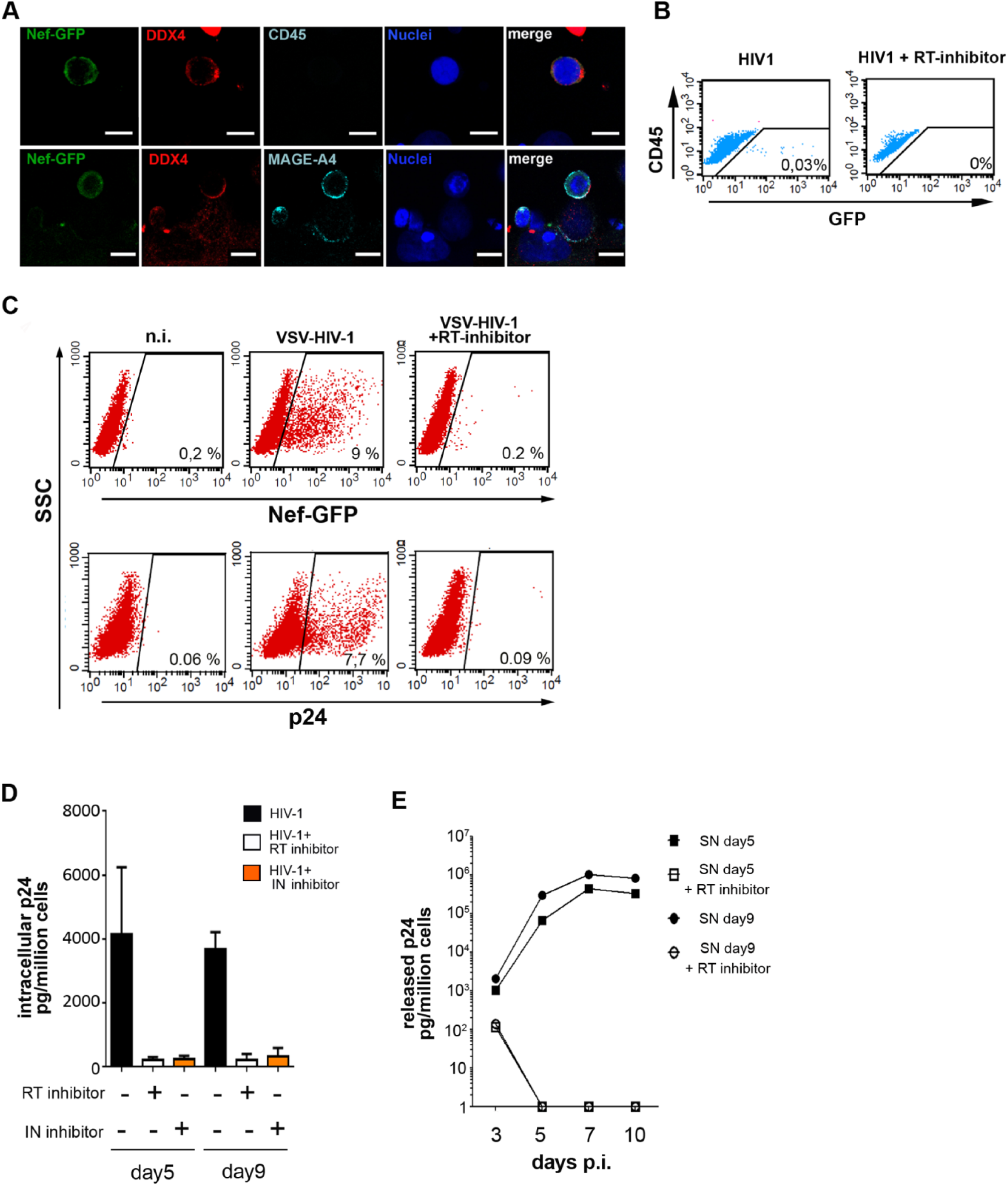
HIV replication in TGCs. (**A, B)** Early HIV-1 replication in germ cells following contact with Jurkat cells infected with HIV-1-*nef*-ires-GFP. **(A)** TGCs incubated with infected Jurkat cells were purified using magnetic beads selection for CD45 and cultured for 9 to 12 days before co-labelling for DDX4 and CD45 (**A**, upper panel) or DDX4 and MAGE-A4 markers (**A**, lower panel). Confocal images show DDX4+ germ cells and MAGEA+DDX4+ early germ cells expressing Nef-GFP. Nuclei were stained with DAPI (blue). Scale bars = 10µm. (**B**) Tcam-2 cells incubated for 18h with Jurkat infected cells and further cultured for 9 days with or without nevirapine. Cells were stained with CD45-PE to discriminate Tcam-2 cells (CD45 negative) from residual Jurkat cells (CD45 positive) and analyzed by flow cytometry for nef-gfp expression. Number of gfp positive events are indicated in the plot. (**C-E**) HIV-1 replication after Tcam-2 exposure to VSV-G-pseudotyped HIV-1. Viral proteins nef-GFP and p24 expression were detected by flow cytometry in Tcam-2 cells cultured for 5 days with or without nevirapine (**C**). Viral production in infected T-cam2 cells cultured for 5 and 9 days with or without RT or integrase inhibitor (raltegravir) was assessed by measuring intracellular p24 in ELISA (**D**). Infectious viral particles release was assessed by exposing PBMCs for 4h to T-cam2 cell supernatants collected at day 5 and 9 post-infection. p24 was measured by ELISA in supernatants from exposed PBMC at the indicated time points (**E**). Data shown are representative of 3 independent experiments.

### Spermatogonia are enriched in HIV early co-factors compared with testicular somatic cells

To further evaluate the permissiveness of TGCs to HIV replication post-entry, we compared the expression profiles of 335 factors that promote or inhibit HIV-1 life cycle post-entry among spermatogonia (SPG), spermatocytes (SPC) and spermatids (SPT) with that in two distinct testicular somatic cell types: testicular macrophages (TM), which are permissive to HIV, and Sertoli cells (SC), which are not a target for HIV, using single cell RNA-sequencing data [42] (Table S1). As expected for an immune cell type, viral sensors (p= 7.76e^-6^) and early inhibitors (p= 1.45e^-2^) were over represented in TM. This is in contrast to TGCs, among which sensors and early inhibitors were even depleted in SPC (p=6.26e^-3^ and 2.59e^-2^, respectively), whereas SC showed an intermediate profile in between that of TGCs and TM (Fig. 7). SPG were significantly enriched in late inhibitors (p= 2.03e^-2^) although less so than TM (p= 1.33e^-8^), whereas SPC (p=1.95e^-4^) and SPT (p= 2.38e^-5^) were depleted (Fig. 7). Interestingly, early co-factors spanning reverse transcription up to DNA integration steps were over-represented in SPG (p= 3.80e^-4^) but not in TM and SC, and tended to be depleted in SPT (albeit not significantly), suggesting a gradient of reduced permissiveness for HIV early replication steps as TGCs differentiate. The distribution frequency of late co-factors (most of them implicated in viral transcription) was comparable for all cell types (Fig. 7). Altogether, these data indicate that TGCs have overall lower innate sensing equipment and low proportion of early HIV inhibitors when compared with SC and TM. Due to their specific enrichment in early co-factors, SPG may represent a more permissive environment for HIV post-entry early steps up to viral DNA integration than late germ cells. Nevertheless, SPG are also enriched in late inhibitors, which could explain low viral production.

**Fig. 7.**
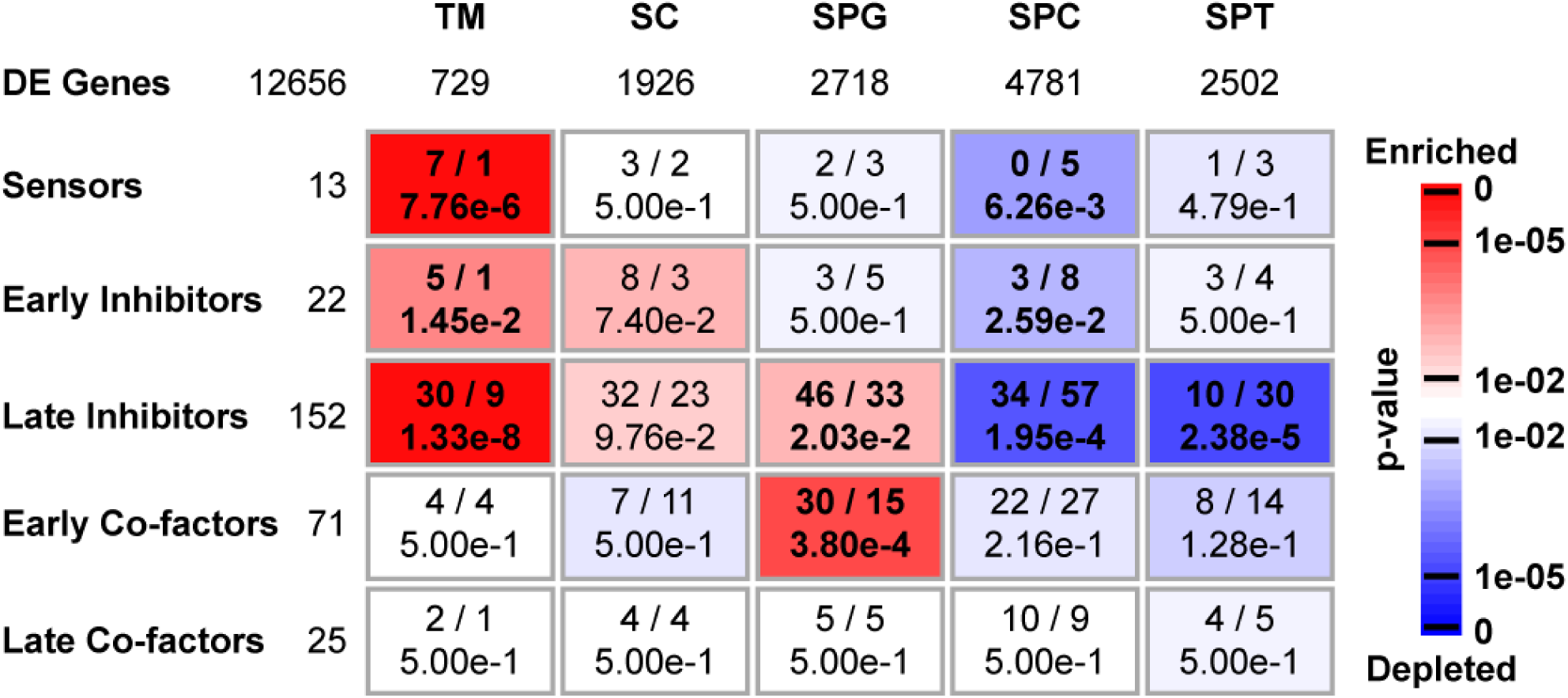
Enrichment analysis of factors involved in HIV life cycle in the human testis. Previously published single-cell RNA-sequencing data of human adult testis [84] were used to identify differentially-expressed (DE) genes showing peak expression in testicular macrophages (TM, 729 genes), Sertoli cells (SC, 1,926 genes), spermatogonia (SPG, 2,718 genes), spermatocytes (SPC, 4,781 genes) or spermatids (SPT, 2,502 genes), including 283 factors involved in HIV life cycle (out of an initial set of 335 factors retrieved from the literature). The over- or under-representation of these factors considered according to different categories (“Early Co-factors”, “Late Co-factors”, “Sensors”, “Early Inhibitors” and “Late Inhibitors”) was then evaluated in each expression cluster by means of a hypergeometric test. Rectangles indicate the observed (left) and expected (right) numbers of genes for each type of factors in each expression cluster, as well as the corresponding p-values adjusted by the Benjamini-Hochberg procedure. Enrichments and depletions are indicated in red and blue, respectively, according to the scale bar.

## Discussion

Here we revealed that primary testicular germ cells can support HIV-1 genome integration and early viral proteins production upon contact with infected lymphocytes *in vitro*. In contrast, cell free virus infection did not lead to detectable viral entry despite HIV-1 binding to TGC and alternative receptors expression. DNAscope on the testis of one infected human and non-human primates demonstrated the presence of isolated HIV or SIV DNA+ TGCs. Using the stem spermatogonia cell line T-cam2, we confirmed that viral integration occurred following cell-associated infection or after bypassing HIV-1 entry, and showed that these early germ cells can produce low level of infectious virions. Gene expression analysis of an array of factors affecting the viral life cycle showed that TGCs were not enriched in defense factors compared with testicular macrophages and Sertoli cells, and that spermatogonia were specifically enriched in co-factors involved in HIV post-entry life cycle up to integration.

While a number of viruses use ubiquitous receptors to attach and enter the cells, HIV virions require specific cellular receptors for infection. Although the main HIV receptor CD4 was absent, TGCs expressed on their surface CD206, GalCer and HSPGs, all previously showed to allow binding, entry or capture of a range of viruses including HIV in various cell types [42– 49]. Ligands of HSPGs, and to a lesser extent of CD206, inhibited binding of cell free virions to TGCs, whereas despite its broad expression, GalCer was not involved, possibly because the detected isoform(s) were unable to bind gp120 [50]. HIV binding to TGC only partly involved HIV envelope gp120, which could be due to HIV envelope-independent uptake by HSPGs, as previously described on other cells [51]. These findings are comparable to that in spermatozoa [15,43–45] and indicate that these receptors for viruses are retained on TGCs surface as they differentiate into spermatozoa. It implies that early TGCs located below the Sertoli cell tight junctions could act as Trojan horse, enabling the virions attached to their surface to cross the blood testis barrier as these cells progress to the seminal lumen during their differentiation. Bypassing the blood testis barrier could allow the virus to shelter from immune recognition by antibodies and immune cells, favoring viral reservoir establishment in the testis. It could also facilitate the release of viral particles into semen and their attachment to sperm.

In our experimental system, HIV entry and reverse transcription was not detected in TGCs exposed to cell-free virus. In CD4-negative somatic cells, free viral particles entry has been reported *in vitro* but was overall inefficient, and viral internalization by vesicular uptake was essentially a dead end with respect to productive infection [46,52–59]. Despite free virus entry limitation, HIV-1 nucleic acids have been evidenced in HIV+ individuals *in vivo* in specific CD4-negative cell types such as renal epithelial cells [60]. Although the mechanism for HIV entry into these CD4 negative cells *in vivo* remains elusive, studies have suggested it may rely on cell-associated infection [54,59,61–63]. Indeed, cell-to-cell contact between infected and non-infected target cells mediate the transfer of HIV-1 to recipient cells with much greater efficiency (100 to 1,000 times) than direct exposure of target cells to cell-free virus [64] and led to viral replication in a few CD4-negative cells, e.g. renal epithelial cells and astrocytes [54,59,63]. In the testis of humans and non-human primates, T lymphocytes and interstitial macrophages infected with HIV/SIV are in close proximity to spermatogonia [28,30,36,37], which are located at the basal compartment of the seminiferous epithelium and therefore not segregated by the blood/Sertoli cell testis barrier. Here we showed that co-culture of infected T cells and TGCs, as well as bypassing HIV entry steps with VSV envelope-pseudotyped HIV-1, led to HIV DNA integration into testicular germ cells’genome. Indeed, integrated viral DNA was measured in TGCs by Alu-gag PCR and was inhibited by a RT inhibitor, demonstrating the specificity of this detection. In addition, early viral protein synthesis was observed in labelled germ cells using confocal microscopy, an indicator of viral integration since unintegrated HIV-1 DNA is associated with transcriptional silencing [65]. Finally, viral proteins detection in TGCs by flow cytometry was inhibited by both RT and integrase inhibitors.

Using next generation *in situ* hybridization, we aimed to determine whether TGCs are infected by HIV *in vivo*. Our results demonstrate the presence of isolated DDX4+ testicular germ cells harboring HIV/SIV DNA in human, as well as in an experimental simian model of HIV infection (rhesus macaque) and a natural host for SIV (African Green monkey), indicating that the virus can proceed early replication steps in both human and simian TGCs *in vivo*. Due to the rarity of testis samples from infected individuals, we could not establish the frequency of HIV DNA detection in TGCs *in vivo*. However our *in vitro* results on TGCs together with our analysis of the testis from an HIV+ donor suggest it is likely to be low.

The detection of HIV/SIV within differentiating germ cells distant from the base of the seminiferous tubules *in situ* could potentially result from clonal infection of early TGCs. Importantly, our data indicate that early TGCs remained viable in culture after viral entry, integration and protein expression, suggesting that viral integration was not detrimental to these cells. Clonal infection of the male germ line up to spermatozoa is suggested by reports of HIV DNA in a low percentage of ejaculated sperm [16–21,23]. Indeed spermatozoa do not support HIV entry, are transcriptionally silent and have a highly compact chromatin, thus preventing HIV replication steps. Patients’ spermatozoa harboring viral DNA *in vivo* were reported to fertilize hamster ova and transfer HIV genome to early embryo [22], indicating that HIV did not impair their fertilization capacities.

The efficiency of HIV replication within a given cell type depends upon a fine balance between factors that positively (co-factors) or adversely (inhibitors) impact the virus life cycle, some of which are actively counteracted by HIV-1 proteins (e.g. APOBEC3G, BST2, SERINC5/3) [66]. Using single-cell transcriptomic data of human testis, we analyzed the expression levels of 335 transcripts that affect HIV replication post-entry in isolated testicular germ cell populations and compared it with that in two somatic testicular cell types, an immune cell type infected by HIV *in vivo* (testicular macrophages) and a non-immune cell type that is not infected by HIV (Sertoli cells) [27]. Although we screened for a large number of known factors, we acknowledge that this analysis is not exhaustive and restricted to transcripts. We found that TGCs were not enriched in viral sensors or early inhibitors. In spermatogonia, except the vRNA sensor RIG-1 and its adaptor IRF3, genes encoding viral RNA, DNA or protein sensors were either not expressed (e.g. TLRs) or their necessary co-factors missing (e.g. IRF7, MyD88, STING), indicating restricted sensing equipment. Transcripts for a range of early HIV inhibitors including IFITM2 and 3 (fusion), PSG-L1 (reverse-transcription), OAS (uncoating) and Mx2 (nuclear import) were also reduced in TGCs versus Sertoli cells and/or testicular macrophages. However, TRIM28 was well-expressed in TGCs compared with both Sertoli cells and testicular macrophages, which could explain limited HIV integration (Table S1). Interestingly, we recently demonstrated that human testicular germ cells were productively infected by Zika virus and that the infection did not impact their survival [26]. In fact, testis antiviral responses were weak [26] and infected TGCs persisted in semen for prolonged duration (article in preparation). We and others also showed that rodent germ cells exposed to a range of viral stimuli did not produce interferon-stimulated antiviral proteins (reviewed in [1]). Altogether, these results suggest that aside their naturally protective testis environment (eg blood testis barrier and neighboring somatic cell types), testicular germ cells do not have strong canonical anti-viral equipment. Further studies are needed to decipher how TGCs respond to viral infections and whether they have evolved specific alternative protective mechanisms to restrict viral replication and its potentially deleterious effects.

Interestingly, early HIV co-factors spanning from reverse-transcription (e.g. DHX9 UBE2B and TRIAD3) to integration steps (LEDGF, INI1, NUP153) were observed at higher frequency in spermatogonia as compared with testicular macrophages, Sertoli cells and late TGCs. We hypothesize that mitotic spermatogonia may therefore be more sensitive to HIV infection than late TGCs, all the more because their localization at the base of the seminiferous tubules could favor cell-to-cell infection by infected leukocytes, thus bypassing entry restriction. Because of their limited life span in culture, we could not determine whether spermatocytes and spermatids were less permissive to HIV replication than spermatogonia *in vitro*.

T-cam2 spermatogonia supported low level of HIV-1 replication up to the production of infectious viral particles, as demonstrated by the infection of PBMCs with infected T-cam2 supernatants and by the inhibition of T-cam2 virus release by RT and integrase inhibitors. *In vivo*, viral RNA and proteins have been detected in a few TGCs within the testis of SIV-infected macaques [36,37], suggesting viral production in a restricted number of cells. Our transcriptomic data analysis revealed that spermatogonia are enriched in a wide range of viral transcription inhibitors (e.g. HDAC1 and 2, negative elongation factor SPT5 and NELF-A/B/CD), which could restrict late viral replication and hence virions production. Altogether, our results support the consensus that male gametes are at best minor contributors to HIV horizontal transmission in comparison with productively infected leukocytes and free viral particles present in semen [35,67,68].

In conclusion, our study reveals that testicular early germ cells have the potential to support low level of HIV integration upon cell-associated infection. Such infection could represent a way for HIV to integrate the germline and become endogenous in the future, as happened during human evolution for a number of retroviruses. Further studies are needed to determine the probability of this event.

## Materials and methods

### Primary testicular germ cells (TGCs) isolation and culture

Normal testes were obtained at autopsy or after orchidectomy from prostate cancer patients who had not received any hormone therapy. The procedure was approved by the National agency for biomedical research (authorization #PF S09-015) and CPP Ouest V local ethic committee (authorization #DC-2016-2783). Only testes displaying normal spermatogenesis, as assessed by transillumination, were used in this study. For receptor detection and HIV binding assay, testes were dissociated with tweezer, incubated in digesting medium (0.5 mg/mL collagenase I, 75ug/mL DNase, 1ug/mL SBTI, PBS 1X) for 90min at 34°C under agitation (110 rpm) and then filtered (100µm). Seminiferous tubules were then mechanically dissociated to avoid trypsin usage, before cell filtration through 300µm and 100µm meshes in PBS containing DNase (40µg/mL). For testicular cells culture and entry assays, testicular tissues fragments were first incubated in digesting medium (2mg/mL hyaluronidase, 2mg/mL collagenase I, 20µg/mL in DMEM/F12) for 60 min at 37°C under agitation (110 rpm). After centrifugation, cell pellet was submitted to trypsin digestion (0.25%, 5mL/g, 20min at 37°C) before trypsin inactivation with FCS, then filtered (60µm), washed 3 times and cultured overnight in DMEM/F12 medium supplemented with 1X nonessential amino acids, 1X ITS (human insulin, human transferrin, and sodium selenite), 100U/mL penicillin, 100µg/mL streptomycin, 10%FCS (all reagents were from Sigma-Aldrich). Floating primary testicular germ cells were collected and cultured onto laminin (Sigma-Aldrich) treated dishes at a density of 20,000 to 40,000 cells/cm^2^ in supplemented StemPro-34 (Invitrogen) as described elsewhere (Sadri-Ardekani et al, JAMA 2009).

### Other cells and testis tissues

Tcam-2 seminoma cell line [69](kindly given by Dr Janet Shipley, The Institute of Cancer Research, London), and CCR5-expressing Jurkat [70] (provided by C. Goujon) were cultured in RPMI-1640 medium. Peripheral blood mononuclear cells (PBMCs) were prepared and cultured as previously described [71]. 293T and positive control cells NP2-CD4, NP2-CCR5, NP2-CXCR4, GHOST-CCR3 and HT-29 cells were maintained in Dulbecco’s Modified Eagles Medium (DMEM) supplemented with 10% fetal calf serum (FCS). All media were supplemented with 100U/mL penicillin, 100µg/mL streptomycin, 2mM L-glutamine and 10% FCS (all reagents from Sigma-Aldrich). Spermatozoa were obtained from healthy fertile volunteer donors with their informed consent (CPP Ouest V local ethic committee authorization #DC-2016-2783) using PureSperm gradient as instructed by manufacturer (Nidacon Laboratories AB, Gotheburg, Sweden). Human testicular tissue was obtained at autopsy from an HIV-1 infected man (donor 108, blood viral load at death 51 680 copies/mL) [72]. NHP testicular tissues were collected at autopsy from an SIVagm+ African green monkey (*Chlorocebus sabaeus*) euthanized 64 day post-infection for another study (blood viral load of 12 085 copies/mL) [73], and from a SIVmac251 Rhesus macaque euthanized at 100 day post-infection (blood viral load > 10^6^ copies/mL) [74].

### Viruses and DNA constructs

Wild type HIV-1 R5 macrophage-tropic (SF162, JR-CSF, BA-L) or X4 strains (IIIb) and primary isolates R5 non-macrophage tropic (ES X-2556-3, subtype B) or R5×4 (ESP-2196-2, subtype G) were obtained from the NIBSC (National Institute for Biological Standards and Control Centralised Facility for AIDS Reagents) and viral stocks produced on PBMCs as previously described [71]. The following viruses were generated by transiently transfecting 293T cells with Lipofectamine 2000 (Invitrogen): 1) *env* deleted or HIV-1 env expressing pseudoviruses using transfer vector pHRsin-cppt-SEW [75], packaging pCMV8.2 [76], HxB2 env pSVIII [77] (kindly provided by Stuart Neil, King’s college, London, UK); 2) HIV-1^GFP-Vpr^ viral particles, using pSG3Δenv, pcDNA3.1D/V5-His-TOPO-env_SF162_, vpr-gfp [78](a kind gift from Christiane Moog, University of Strasbourg, France); 3) VSV G-pseudotyped HIV-1 using full-length HIV-1 molecular clone pNL4-3 [79] or the R5-tropic pNL4-3 AD8 derivative [80] and VSV-G envelope encoding plasmid (PT3343-5, Clontech); 4) VSV G-pseudotyped HIV-1 *nef*-ires-GFP using pBR_NL4-3 IRES-eGFP (X4-tropic) or pBR_NL4-3 92TH014.12 IRES eGFP (R5 tropic) [81] (a kind gift from Frank Kirchhoff, Ulm University, Germany) and VSV-G envelope encoding plasmid (PT3343-5, Clontech). 293T cell supernatants were collected 3 days post-transfection, filtered using 0.45 µm membrane and stored at -80°C.

### Flow cytometry for antigen detection and cell sorting

Immunostaining of surface antigens was performed as previously described [82]. The following antibodies were used: anti-CD45 (-PE or PC7 conjugated, HI30, BD Pharmingen), anti-HLA-ABC-PE (G46-2.6, BD Pharmingen) anti-Vimentin (EPR3776, 1µg/mL, Epitomics), anti-DDX4 (Rabbit polyclonal, 5µg/mL, Abcam), anti-MAGEA4 (clone 57B, 4µg/mL) [39], anti-CD4 (ARP337, 10µg/mL), anti-CXCR4 (ARP3101 12G5, 10µg/mL) (both from EVA/MRC, NIBSC), anti-CCR5 (45549, 10µg/mL, R&D), anti-CCR3 (61828, 5µg/mL, R&D), anti-GalactosylCeramide (mGalC, 10µg/mL, Millipore), anti-Heparan Sulfate-ProteoGlycans (10E4, 5µg/mL, Seikagaku corporation) and anti-p24 gag RD1 (KC57, Beckman coulter). PE conjugated anti-mouse or anti-rabbit Ig (Jackson Immunoresearch) were used as secondary antibodies. Control isotype antibodies were used as negative controls, except for HSPGs where cells were pretreated or not with pronase (3,5mg/mL, Roche) and then subjected to immunostaining. The expression of Mannose Receptor (MR) was analyzed as previously described [15]. Cellular DNA content was determined using DRAQ5 (20uM, Biostatus Limited). FACScalibur flow cytometer (Becton Dickinson, Franklin Lakes, USA) and CELLQuestPro Software were used for acquisition and analysis.

For cell sorting, TGCs were stained with the fixable viability stain FVS450 (BD Biosciences), as recommended by the supplier before fixation in 1,5% paraformaldehyde. Cells were then labelled using DDX4 anti-rabbit-PE antibodies, CD45-PC7 Ab (clone J33, 1/15, Beckman Coulter), CD3-BV510 Ab (clone HIT3a, 1µg/mL, BD Biosciences), HLA-DR-BV510 Ab (clone G46-6, 7µg/mL, BD Biosciences) and MAGEA4 Ab (clone 57B, 4µg/mL) [39] previously coupled to Alexa Fluor 647 using Zenon labeling kit (Molecular Probes). Matched isotype antibodies were used as negative controls. Cells negative for CD45, CD3, HLA-DR and positive for DDX4 and/or MAGEA4 were sorted using FACSAria flow cytometer (Becton Dickinson, Franklin Lakes, USA) connected to the DIVA software. For Tcam-2, CD45-PC7 antibody was used for CD45 negative T-cam2 cell sorting.

### Binding assay

The binding assay was performed as previously described [15], with some modifications. Briefly, cells were incubated with indicated HIV-1 strains (25 or 75 ng p24 per 10^6^ cells) in a final volume of 150µL of medium and incubated for 2 hours at 37°C in RPMI-1640 2,5% FCS. Cells were then thoroughly washed, lysed and the p24 content determined by ELISA. When indicated, the assay was performed on cells exposed to Pronase (3,5mg/ml) or using virus pre-exposed to 1 to 100 U/mL heparin (Sigma-Aldrich). The role of mannose receptor was evaluated by incubating cells with HIV-1 in the presence of 100µg/mL BMA (Sigma-Aldrich) or 100µg/mL bovine serum albumin (BSA) as control. Neutralization assays were performed by pre-incubating cells with anti-GalactosylCeramide antibody (mGalC, 10µg/mL, Millipore 10µg/mL, 30 min at 4°C) or viral inoculum with monoclonal antibody against HIV-1 gp120 (1-2µg/mL, clone F105, AIDS Reagents, NIBSC, 30 min at 37°C). Matched isotype antibodies were used as negative controls. Controls for nonspecific viral attachment were systematically performed by incubating viral inoculum in the absence of cells. In all experiments, each condition was performed in duplicate and residual p24 level deduced.

### Cell infection

TGCs, Tcam-2 and Jurkat cells were infected with 200ng p24/10^6^ cells of the indicated virus as previously described [82] and cultured with or without 37,5µM nevirapin (non-nucleosidic reverse-transcriptase inhibitor) or 100nM raltegravir (integrase inhibitor) (both from AIDS Reagents, NIBSC), as specified. For cell-associated infection, Jurkat donor cells were infected with the indicated VSV-G pseudotyped HIV NL4-3 strain. TGCs or T-cam2 were incubated for 18h with infected Jurkat cells (40 to 80% positive for HIV-1 Gag or GFP). Donor cells were then removed by positive selection of CD45+ cells (CD45 microbeads kit, Miltenyi Biotec) (TGCs) or pipetting (Tcam-2) and target cells put in culture. GFP content was analysed at the indicated time points in DDX4 positive cells by confocal microscopy (TGCs) or by flow cytometry in CD45 negative Tcam-2 cells. For integrated viral DNA quantification, TGCs and Tcam-2 were further purified by FACS sorting as described above (see Flow cytometry and cell sorting).

### Confocal microscopy for viral entry

Germ cells were pre-labelled with 10uM CFSE or CellTrace Violet (Invitrogen) before being exposed to free viral particles overnight and treated with 0.25% trypsin-EDTA (Sigma-Aldrich) for 5 min at 37°C post-infection to remove virions attached to the cell surface. Cells were loaded onto polylysine-coated coverslips, fixed in 4% para-formaldehyde, washed in PBS and incubated in a blocking/permeabilizing solution (0.3M Glycine, 0.05%Triton-X, 3% BSA, 1X PBS) for 30 min at RT. Cells were then stained with the following antibodies: mouse monoclonal anti-p24 (183-H12-5C, EVA, NIBSC), rabbit anti-DDX4 (5µg/mL, Abcam), mouse anti-MageA4 (clone 57B, 4µg/mL) [39], CD45-Alexa-fluor 647 (J.33 clone, Beckman coulter). Anti-mouse Alexa-fluor 555 (Thermo Fisher Scientific), anti-rabbit Alexa-fluor 488 (Thermo Fisher Scientific) and Alexa-fluor 647 (Jackson Immunoresearch) were used as secondary antibodies. Isotype control antibodies or mock infected cells were used as controls. Nuclei were stained either with 10 μM DRAQ5 (Biostatus) or with DAPI (Molecular Probes), and slides mounted with Prolong Gold Antifade mountant (Molecular Probes). Images were acquired with the SP8 confocal system microscope (objective 60X, Z-step 0.3 to 0.5µm intervals) (Leica) connected to LAS software and analyzed with Fiji software.

### Real-Time PCR for HIV-1 DNA

Intracellular HIV-1 DNA was measured as previously described [72]. Input virus was treated with benzonase (500U/mL, Sigma-Aldrich) before cell exposure to reduce nucleic acid contamination in the viral stock. Cells infected in the presence of the RT inhibitor nevirapin (37,5µM) were used to discriminate DNA originated from input viruses. Integrated HIV DNA was quantified as previously described using Alu-gag PCR assay [83].

### Next generation *in situ* hybridization (DNAscope)

DNAscope was used for the detection of viral DNA for both HIV and SIV, as previously described [74]. To immunophenotype the cells containing vDNA, DNAscope detection by Tyramide Signal Amplification (TSA™) Plus Cy3.5 was combined with immunofluorescence targeting cell markers using rabbit polyclonal anti-DDX4 (1:500; Abcam) for TGC and mouse monoclonal anti-CD4 (1:100; clone 1F6, Vector) for T-lineage cells. 4 to 6 5μm sections were run per sample. An average of 90982mm^2^ of tissue area was screen to find positive event. Fluorescent slides mounted with Prolong^®^ Gold (Invitrogen) were imaged on an Olympus FV10i confocal microscope using a 60x phase contrast oil-immersion objective (NA 1.35) and applying a sequential mode to separately capture the fluorescence from the different fluorochromes at an image resolution of 1024 x 1024 pixels.

### p24 ELISA and infectivity assay

p24 concentrations in T-cam2 cells were measured by ELISA (Innogenetics, according to manufacturer’s instructions) and the infectivity of viral particles determined through PBMCs exposure to T-cam2 cell supernatants for 4 h, with p24 quantification at the indicated time post-infection.

### Analysis of single-cell RNA-sequencing data

#### Data processing

Single-cell RNA-sequencing data of human adult testicular cells [84] were recovered from the ReproGenomics Viewer [85,86] and processed using the AMEN software [87]. Briefly, out of the 23’109 genes measured in this experiment, only those with an average expression level (log2[count+1]) of at least 1 in a least one cell population and a fold-change of at least 2 between at least two cell populations were considered as being differentially-expressed. Selected genes were further classified according to their peak expression in testicular macrophages (Cell population tMϕ in the initial publication of [84]), in Sertoli cells, in spermatogonia (cell populations SSC1, SSC2, differentiating and differentiated SPG in the original publication), in primary or secondary spermatocytes (cell populations L1, L2, L3, Z, P, D and SPC7 in the original publication) or in round or elongated spermatids (Cell population S1 to S4 in the original publication). The consistency of the results was controlled by comparing expression profiles of all selected genes to those obtained by Hermann *et al.* [88] (Fig. S3).

#### Enrichment analysis

A list of known factors involved in HIV life cycle post-entry was retrieved from the literature (Table S1). These were annotated as “Early Co-factors” (79 genes), “Late Co-factors” (28 genes), “Sensors” (15 genes), “Early inhibitors” (28 genes) or “Late inhibitors” (185 genes) according to their demonstrated roles. The over- and under-representation of these factors among clusters of differentially-expressed genes was then evaluated with a hypergeometric test, while the false discovery rate was controlled by adjusting resulting p-values using the Benjamini and Hochberg method [89].

### Statistical analysis

Statistical analyses were performed using commercially available software (GraphPadPrism 6, GraphPad Software, Inc., La Jolla, California, USA). Data were analysed with non-parametric Friedman-Dunn’s test or Mann Withney test, as indicated in figure legends. Values were considered significant when P<0.05. Statistical analyses performed on single-cell RNA-sequencing data are described above in the relevant section.

## ACKNOWLEDGEMENTS

This project received funding from Inserm, ANRS, Sidaction, CAPES-COFECUB and was funded in part with Federal funds from the National Cancer Institute, National Institutes of Health, under Contract No. HHSN261200800001E. Experiments were conducted in part on L3, MRic, H2P2 and flow cytometry platforms at Biosit federative structure (Univ Rennes, CNRS, Inserm, BIOSIT [(Biologie, Santé, Innovation Technologique de Rennes)] -UMS 3480, US_S018). This project has been funded in part with The content of this publication does not necessarily reflect the views or policies of the Department of Health and Human Services, nor does mention of trade names, commercial products, or organizations imply endorsement by the U.S. Government.

## SUPPLEMENTAL INFORMATIONS

**Fig. S1.**
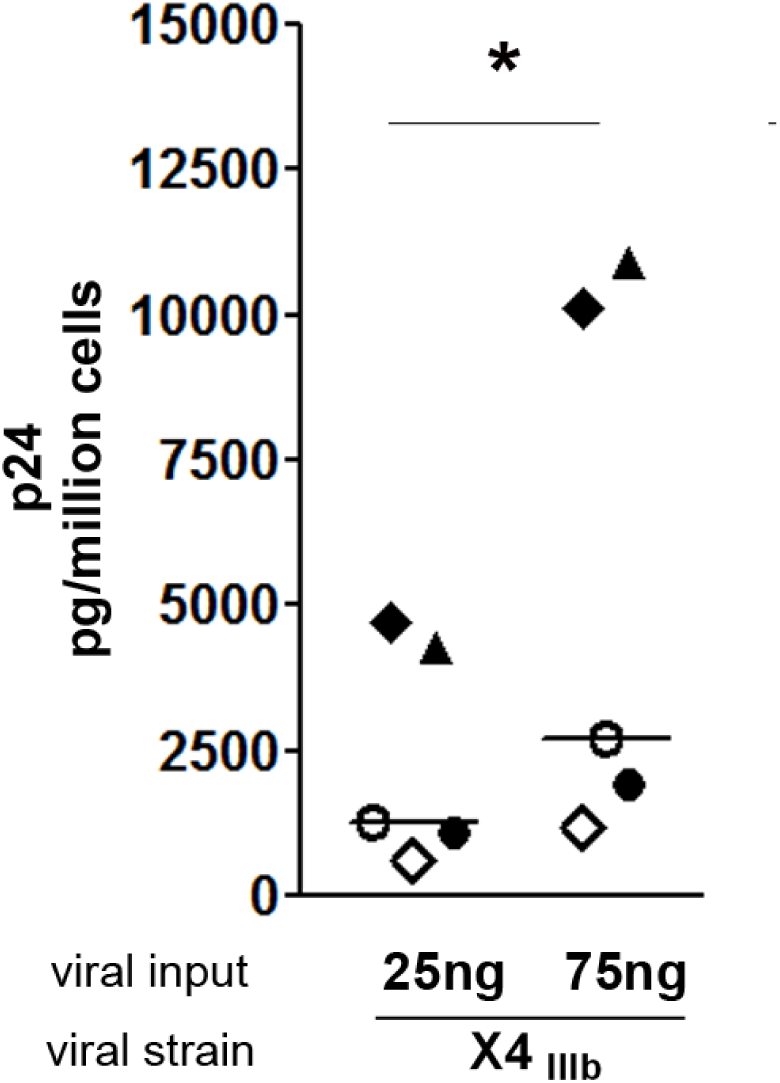
HIV-1 binding of X4_IIIb_ HIV strain to primary TGCs. TGCs isolated from 6 donors were tested for their ability to capture X4_IIIB_ following a 2h incubation at 37°C with the indicated amount of virus, as measured by p24 ELISA on cell lysates following thorough washes and deduction of p24 background measured in wells with no cells. Statistical analysis with non-parametric test: Wilcoxon test. * p< 0.05; ** p< 0.01; *** p< 0.001. Each donor is represented by a specific symbol.

**Fig. S2.**
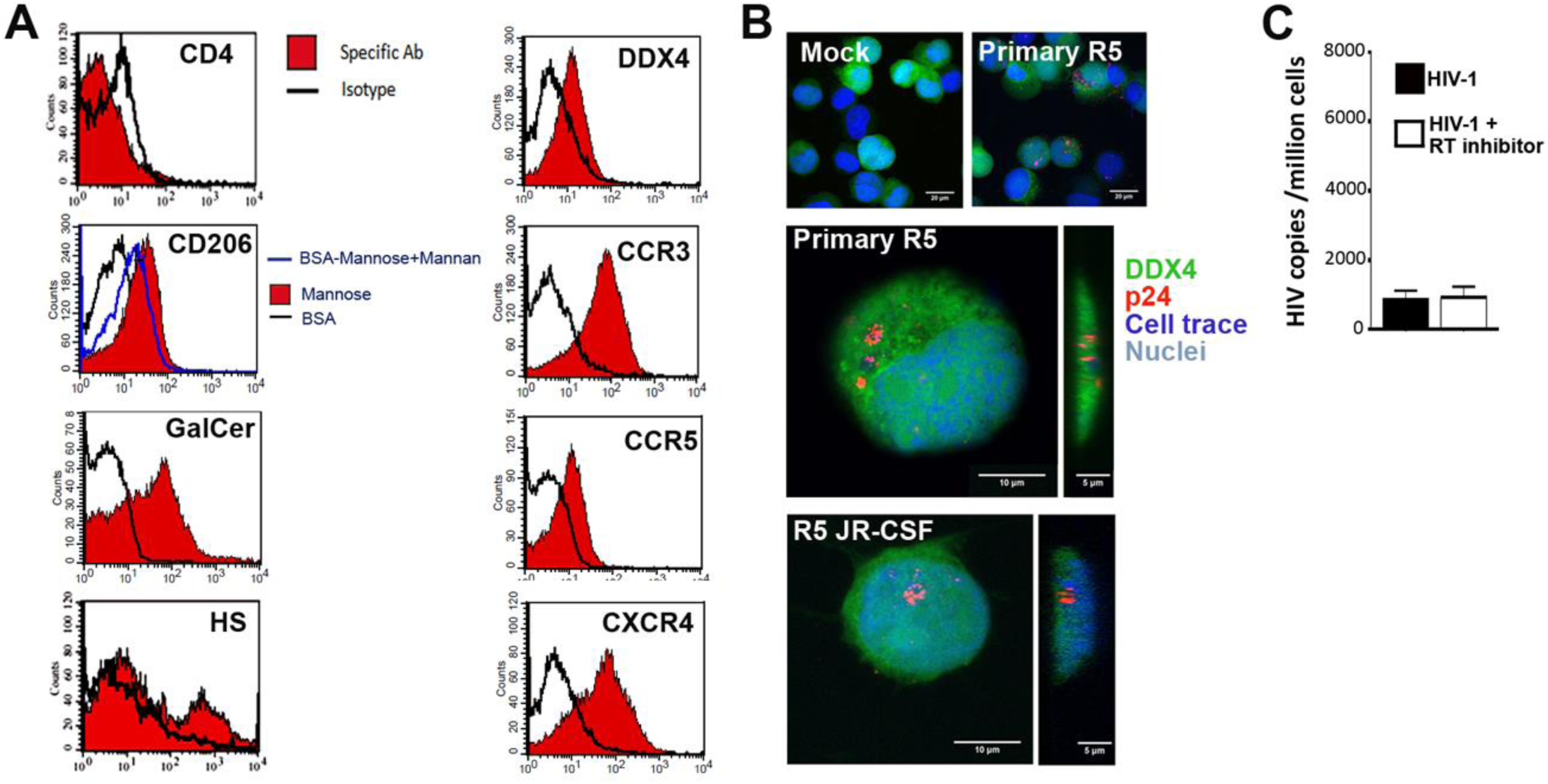
HIV-1 alternative receptors expression and cell-free entry in Tcam-2. (**A**) Potential HIV receptors was assessed on the surface of Tcam-2 cells labelled for DDX4 using specific antibodies against CD4, CCR5, CXCR4, CCR3, GalactosylCeramide (GalCer) and HSPGs, or following detection of BSA-Mannose, a ligand for mannose receptor. Immunolabelling was analyzed by flow cytometry. (**B**) HIV-1 entry was assessed in CFSE-labelled Tcam-2 cells (in green) exposed for 4h to a primary R5-tropic HIV-1 strain (ES X-2556-3) or mock exposed, before immunolabelling with p24 antibody (red). Nuclei were stained with DAPI (blue). (**C**) HIV-1 reverse transcription was assessed by qPCR on Tcam-2 cells cultured 24h with or without the reverse transcriptase inhibitor nevirapin, following exposure to HIV R5_JR-CSF_ HIV-1 (n=6).

**Fig. S3.**
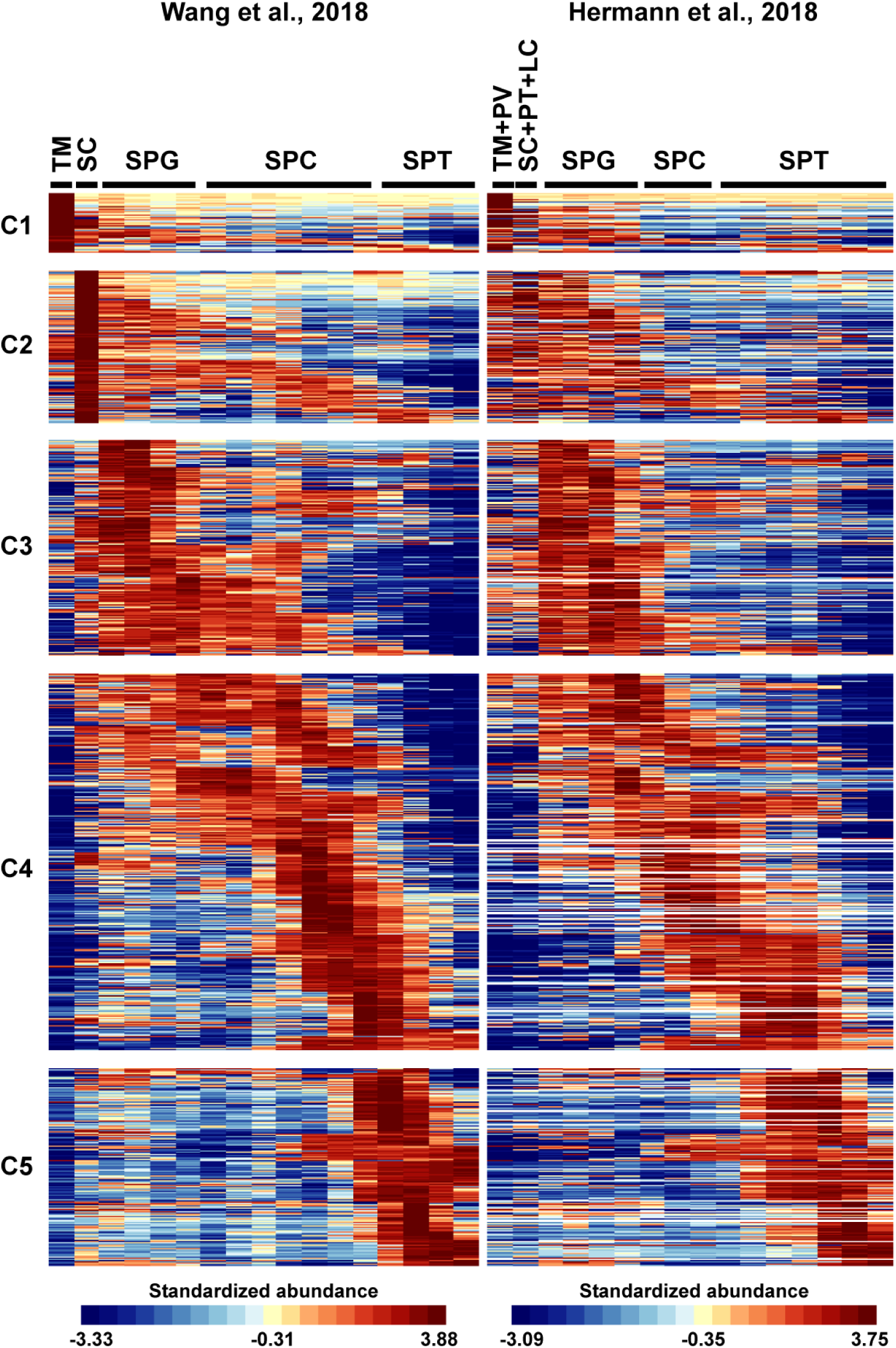
Dynamics of gene expression in human testicular cells as evidenced by single-cell RNA-sequencing. A false-color heatmap representation of gene expression in distinct human testicular cell populations is shown. Briefly, differentially-expressed genes were identified according to single-cell RNA-sequencing data of human testis from Wang and colleagues [84] and grouped into 5 expression clusters (C1 to C5) according to peak expression in testicular macrophages (TM), Sertoli cells (SC) spermatogonia (SPG), spermatocytes (SPC) or spermatids (SPT), respectively. Corresponding expression profiles from the single-cell RNA-sequencing dataset of Hermann and collaborators [88] are displayed on the right. Each line is a gene and each column is a cell type. Standardized abundances are displayed according to the scale bars for each dataset. TM+PV = Testicular macrophages and Perivascular cells; SC+PT+LC = Sertoli cells, peritubular cells and Leydig cells.

**Table S1 List of HIV replication modulating factors expressed by human testicular macrophages and germ cells as detected in single-cell RNA-sequencing data.**

A searchable excel file containing various information on 335 factors involved in HIV life cycle is provided, including: the Gene Name (column A), the Entrez Gene ID (column B), the Type of Factor (e.g. “Early inhibitor”; Column C), the Step/Process they affect (e.g. “Nuclear import”; Column D) and the corresponding Pubmed ID(s) (Column E). Subsequent column indicate the Expression Cluster to which they belong (e.g. C1-Macrophages; Column F), according to single-cell RNA-sequencing data from Wang et al. [84]. Finally, columns G to V provide the corresponding average gene expression values (log-2 transformed) in each testicular cell population and are color-coded according to low (blue) or high (red) expression levels.

